# Local Extensional Flows Induce Long-Range Fiber Alignment in 3D Collagen Hydrogels

**DOI:** 10.1101/2022.02.04.479166

**Authors:** Adeel Ahmed, Indranil M. Joshi, Mehran Mansouri, Ann M. Byerley, Steven W. Day, Thomas R. Gaborski, Vinay V. Abhyankar

**Affiliations:** Biomedical Engineering, Kate Gleason College of Engineering, Rochester Institute of Technology

**Keywords:** collagen, alignment, microfluidic, 3D hydrogel

## Abstract

Randomly oriented type I collagen (COL1) fibers in the extracellular matrix (ECM) are reorganized by biophysical forces into aligned domains extending several millimeters and with varying degrees of fiber alignment. These aligned fibers can transmit traction forces, guide tumor cell migration, facilitate angiogenesis, and influence tissue morphogenesis. To create aligned COL1 domains in microfluidic cell culture models, shear flows have been used to align thin COL1 matrices (<50μm in height) in a microchannel. However, there has been limited investigation into the role of shear flows in aligning 3D hydrogels (>130μm). Here, we show that pure shear flows do not induce fiber alignment in 3D atelo COL1 hydrogels, but the simple addition of local extensional flow promotes alignment that is maintained across several millimeters, with a degree of alignment directly related to the extensional strain rate. We further advance experimental capabilities by addressing the practical challenge of accessing a 3D hydrogel formed within a microchannel by introducing a magnetically coupled modular platform that can be released to expose the microengineered hydrogel. We demonstrate the platform’s capability to pattern cells and fabricate multi-layered COL1 matrices using layer-by-layer fabrication and specialized modules. Our approach provides an easy-to-use fabrication method to achieve advanced hydrogel microengineering capabilities that combine fiber alignment with biofabrication capabilities.

## 1. Introduction

The extracellular matrix is a 3D fibrous network that provides structural integrity to tissues and guides cell behavior^1,2^. Physical properties such as stiffness, fiber density, and fiber organization serve as cues to guide cell motility, morphology, and phenotype^3–5^. In turn, the fiber organization of the ECM is dynamically remodeled by cells in response to local biochemical and biophysical stimuli. These cell-ECM interactions can reorganize randomly oriented collagen type I (COL1) fibers into aligned fiber domains with varying degrees of alignment^6,7^. Due to the non-linear mechanical properties of fibrous COL1 networks, fiber alignment produced by cell-induced traction forces can extend far beyond the local source to establish long-range mechanical communication through which cell position and orientation information can be efficiently transmitted between cell populations^8–11^. Aligned COL1 fibers also influence physiological processes across several length scales (e.g., millimeter to centimeter), including tumor cell invasion^6^, immune cell homing^12^, capillary formation^13–15^, muscle cell orientation^16,17^, and tissue morphogenesis^18^. Given the importance of COL1 fiber alignment in native tissue, several in vitro techniques have emerged to create aligned COL1 matrices and investigate cellular responses in an experimentally tractable context^19–22^.

Microfluidic approaches have become increasingly popular to establish physiologically relevant in vitro culture models because they can incorporate biomaterials and provide robust control over microenvironmental cues ^23–25^. Lee and co-workers first demonstrated the alignment of COL1 fibers by injecting a neutralized, self-assembling COL1 solution into microchannels up to 100 μm wide and 50 μm tall ^26^. They hypothesized that the channel walls acted as nucleation sites onto which COL1 fibrils attach and grow in the flow direction under the influence of shear forces. This mechanism was further investigated by Saeidi et al., who showed that when a self-assembling COL1 solution flowed over a glass substrate, COL1 fibrils tethered to the surface^27^. The surface-bound fibrils experienced a stretching force due to fluid-induced shear at the walls and produced directional growth of COL1 fibers. Other studies have also used shear flows in microfluidic channels to fabricate thin (<50 μm) layers of COL1 fibers on a glass substrate to study corneal keratocyte migration and alignment of nerve cells ^28–30^. These early studies suggest that a stretching force is required to align the self-assembling COL1 subunits and promote directional fiber formation. Conversely, collagen subunits not tethered to a surface experience a torque in the bulk fluid that rotates the subunits and negatively impacts alignment^27^. As a result, aligned COL1 matrices fabricated using shear flows are generally limited to surface coatings or thin gels < 50 μm in thickness. Although these surface coatings are useful in studying 2D cell responses, they cannot be used to encapsulate cells and replicate 3D cell-matrix interactions found in vivo^31–34^.

Extensional flows, characterized by an increase in fluid velocity along the flow direction (i.e., stretch), are commonly used to promote the self-assembly of polymer melts^39,40^, unfold DNA strands^41,42^, or aggregate proteins^43^. In the context of COL1 fibers, extensional flows have been shown to induce self-assembly of COL1 monomers drawn from a droplet into aligned fibers^44^. Extrusion and bioprinting platforms have also used extension to control fiber alignment in collagen matrices, sheets, and threads^45–48^. In our previous work, we demonstrated that the direction and degree of fiber alignment could be controlled over a sub-millimeter length scale as a function of the extensional strain rate 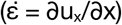 in 60 μm tall microchannels^49^.

To address the growing need for 3D COL1 matrices with ordered fiber domains, techniques such as magnetic fiber alignment^35,36^, external mechanical strain^7,37^, and cell-mediated compaction^38^ have been used to achieve free-standing structures with millimeter-scale alignment. Although useful, these approaches require multi-step protocols and are difficult to integrate into microfluidic systems that provide precise control over biophysical and biochemical microenvironment cues. Here, we introduce a simple approach, using a “segmented” microfluidic channel design that creates local extensional flows to generate 3D COL1 hydrogels (>130 μm thick) with long-range fiber alignment. To build additional experimental functionalities, we use specialized modules to pattern cells and demonstrate the layer-by-layer fabrication of heterogeneous hydrogel constructs.

## 2. Results

### 2.1 Shear flows do not align 3D COL1 hydrogels in a microchannel

Shear flows have been used to fabricate surface coatings (< 50 μm) of aligned COL1 fibers using acid solubilized telocollagen, which are well-suited for 2D cell-culture studies ^26–28^. However, a 3D matrix is a more relevant representation of the in vivo environment and is important to investigate cell responses to biophysical cues in vitro, especially in the context of tumor metastasis, developmental biology, and drug development^33,50^. Enzymatically extracted atelocollagen from bovine sources is now commonly used for bioprinting and 3D culture studies because it is easier to manipulate over extended periods of time due to slower self-assembly kinetics and less sensitivity to environmental conditions^51^. Using this material, we investigated whether shear flows, characterized by the shear rate 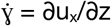 (i.e., the velocity gradient over the channel height), could be used to align 3D atelo COL1 hydrogels.

We injected neutralized, 2.5 mg mL^−1^ COL1 solution into a microfluidic channel with a constant height of 130 μm at shear rates spanning 50 to 1000 s^−1^, and confirmed that there was no extensional component to the flow (i.e., the extensional strain rate, 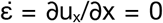) using particle image velocimetry (PIV) **(Figure S1).** After injection, the COL1 solution was allowed to self-assemble in the microchannel in a humidified incubator and imaged using confocal reflectance microscopy (CRM). We measured the fiber coefficient of alignment (CoA), defined as the fraction of fibers oriented within ±15° of the mode value in the fiber angle histogram. As shown in **Figure 1A**, increasing the shear rate did not induce fiber alignment in the 130 μm channels (i.e., CoA < 0.5). Representative images of the COL1 gels are shown in **Figure 1B** and provide visual evidence of unaligned fibers. Based on these results, we concluded that shear flows were not sufficient to align thick atelo COL1 hydrogels, and we explored alternative flow-based methods to create alignment.

**Figure 1:**
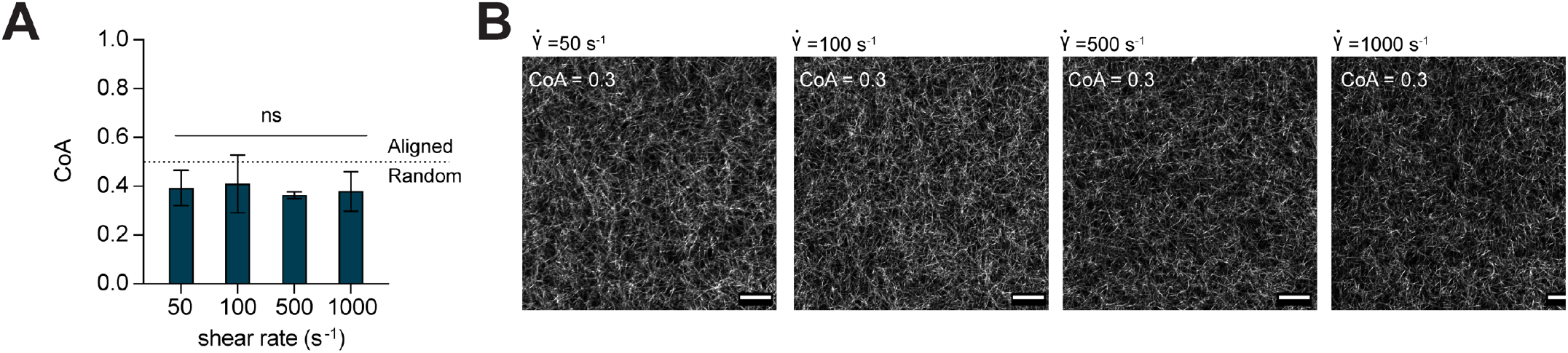
**(A)** Data show COL1 matrices that were formed after injection into straight microchannels at shear rates from 50 - 1000 s^−1^ were not aligned. The dotted line indicates the threshold CoA value for alignment (CoA=0.5). **(B)** Representative CRM images provide visual evidence that the resulting gels were unaligned. Data presented as mean ± SD, n=3, ns=not significant. Scale bar = 25 μm.

### 2.2 A segmented channel design introduces local extensional flows

In our previous work, extensional flow was shown to align 60 μm thick COL1 hydrogels^49^. We also observed that the magnitude of the extensional strain rate directly influenced the degree of fiber alignment. The extensional strain rate has also been identified as a key player in controlling alignment during extrusion and fiber drawing processes in non-microfluidic applications^14,44,45^. Thus, we hypothesized that COL1 fiber alignment could be controlled in 3D hydrogels by exposing self-assembling COL1 solutions to extensional flow in a microchannel.

To introduce extensional flow, we designed a microchannel with five segments, each of which was 5 mm long with a constant height of 130 μm **(Figure 2A)**. The first segment was 10mm wide, every subsequent segment was half the width of the prior segment, and the last segment was 0.75mm wide (labeled a-e in **Figure 2B**). At a constant flow rate Q, we expected the flow velocity to increase at each constriction as the cross-sectional area decreased and then reach a constant value within each segment. To test the approach, neutralized COL1 was injected into the segmented channel at Q = 50μL min^−1^, and the fluid velocity was measured at the constriction ROI (yellow boxes in **Figure 2B**) and within each segment using PIV **(Figure 2B).** The measured extensional strain rate was < 1s^−1^ in constrictions a-b and b-c, increased to 2.7±0.67 s^−1^ in constriction c-d, and reached a maximum value of 9.1±0.52 s^−1^ in constriction d-e. As expected, the extensional strain rate was localized to the constriction region and was approximately zero (< 0.5 s^−1^) in each segment **(Figure S2).**

**Figure 2:**
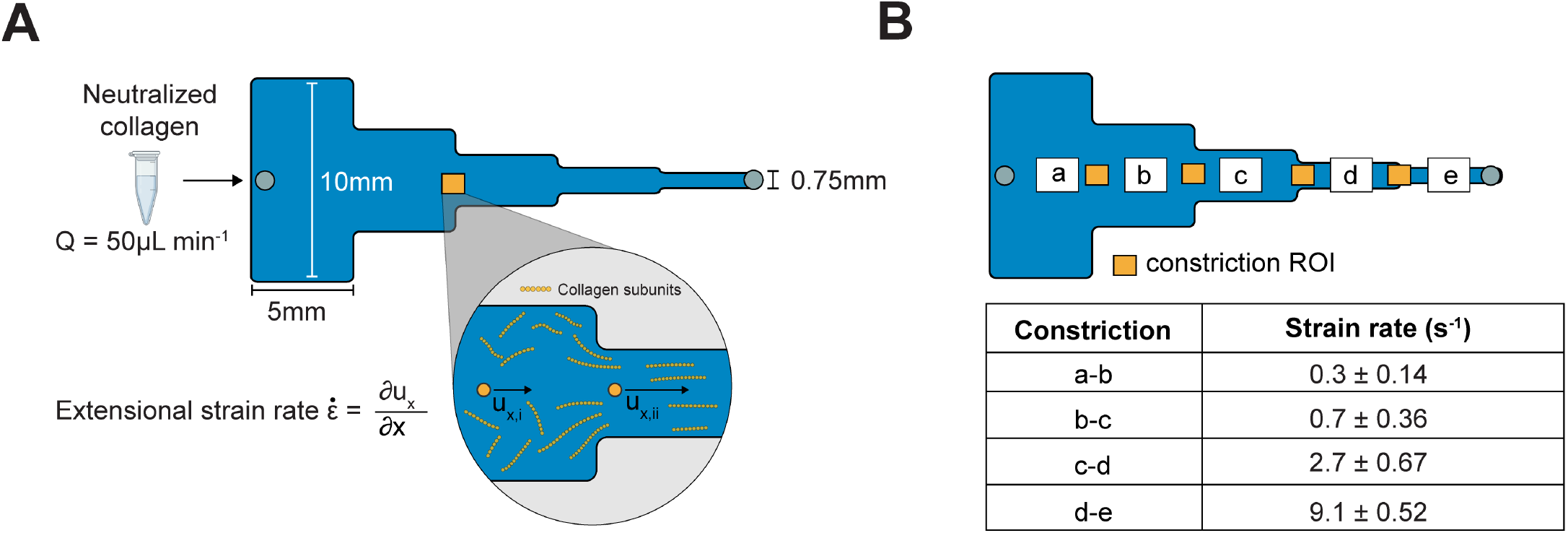
**(A)** Schematic top view of the the 25mm long, segmented channel used to create extensional flow. Each segment is 5mm in length with widths decreasing sequentially from 10mm to 0.75mm left to right. As the cross-sectional area changes, the COL1 subunits experience an extensional flow with a strain rate defined as the change of velocity along the flow direction (left to right) at the entrance of each constriction. **(B)** The extensional strain rate measured by PIV in the constrictions of the segmented channel increases along the direction of flow ranging from 0.3 ± 0.14 s^−1^ to a maximum of 9.1 ± 0.52 s^−1^. Data reported as mean ± SD, n = 3.

After characterizing the extensional flow in our segmented channel, we investigated whether the flow could induce COL1 fiber alignment in 3D gels. We injected neutralized COL1 solutions as previously described and quantified the fiber CoA in each segment as a function of the 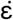 at the constrictions (labeled a-e in **Figure 3A**). Image analysis revealed that when 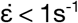 (segments a-c), the mean CoA was below the alignment threshold of 0.5. At higher extensional strain rates, 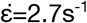 (segment d) and 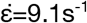 (section e), fibers were aligned with a mean CoA of 0.58±0.13 and 0.64±0.11, respectively **(Figure 3B),** suggesting that the extensional strain rate promoted fiber alignment. The CoA across the different segments showed similar trends when COL1 was injected into the channel at different flow rates (Q = 100, 200, 400, and 1000μL min^−1^) **(Figure S3).** The highest CoA value obtained was CoA=0.8, suggesting an upper limit on the fiber alignment using extensional flows with our current design. Next, we investigated whether extensional flow could align thicker COL1 hydrogels by injecting solutions into a 250μm tall, segmented channel at a flow rate of 750μL min^−1^. The fiber alignment was similar to the 130 μm tall channels (**Figure S4**), with the mean CoA in segment (d) equal to 0.67 ± 0.002 and 0.75 ± 0.01 in segment (e). These data provide evidence that the extensional flow in the segmented channel promoted COL1 fiber alignment in thick hydrogels.

**Figure 3:**
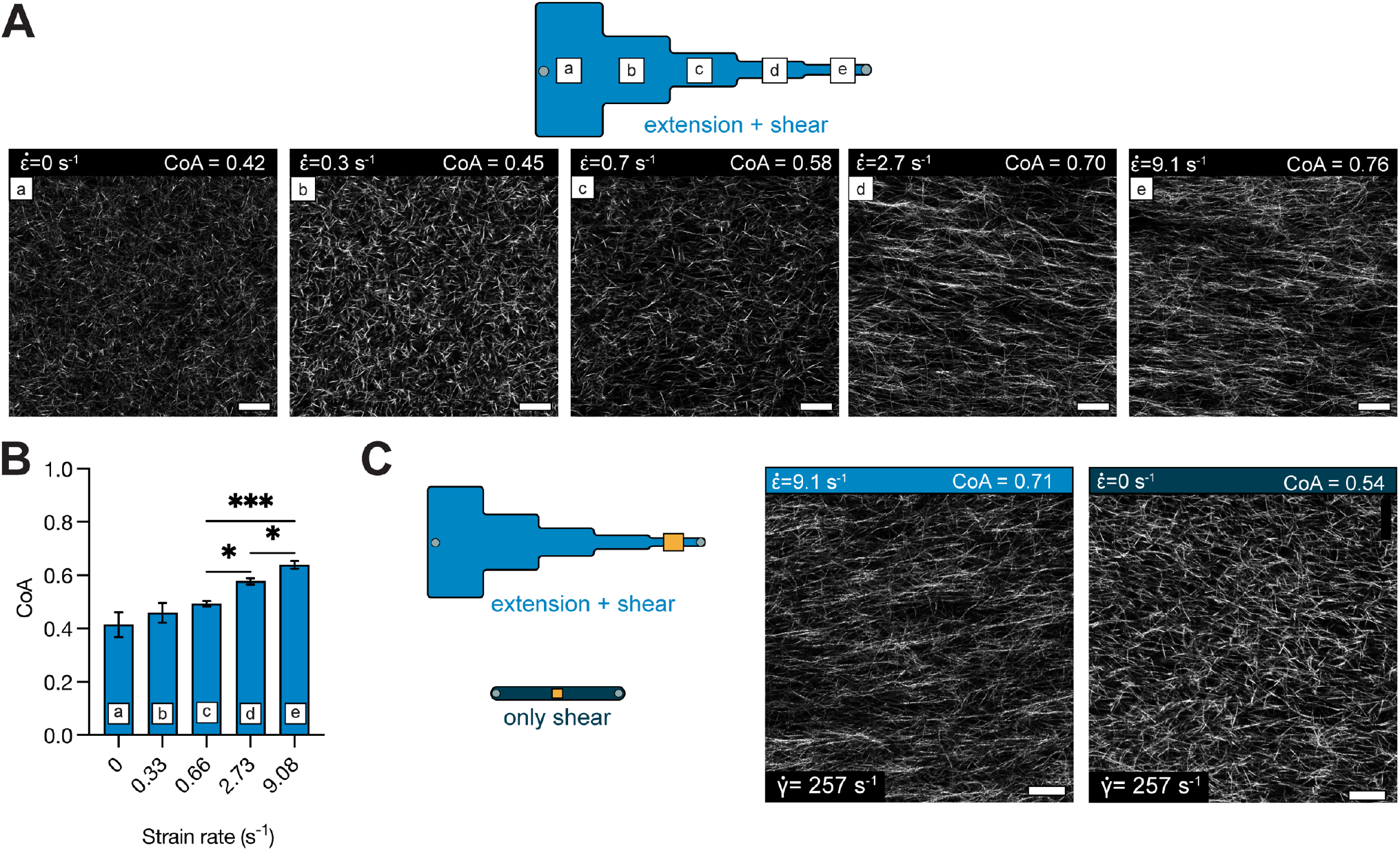
**(A)** Schematic showing the different segments of the segmented channel and the zero-strain channel. Panels (a-e) show representative images of COL1 fibers in the corresponding section of the segmented channel. The degree of alignment of COL1 fibers is seen to increase across the different segments. **(B)** Bar plot showing the mean CoA ± SD in each segment, for COL1 injected at a flow rate of 50μL min^−1^. The CoA was greater than 0.5 at 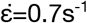. (C) Representative images of COL1 fibers at 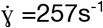 with 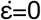 and with 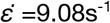. Fiber alignment can be visually confirmed to be higher with extensional strain. data represented as mean ± SD. n = 4. * p<0.5, ** p<0.01

The flow in the segmented microchannel has shear and extensional components, and we sought to understand the role of each on fiber alignment. To decouple the shear rate from the extensional strain rate, we determined the maximum centerline velocity (u_x,max_) in each segment of the segmented channel using PIV and calculated the corresponding shear rate from the equation 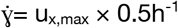 (**Table S5**). The shear rates ranged from 23 s^−1^ in segment (a) to 257 s^−1^ in segment (e). Collagen was then injected into a uniform, straight channel of 130m thickness, such that the shear rate in the uniform channel was matched to the shear rate in a segment, and the extensional component in the uniform channel was ~0 (**Figure S1**). At a shear rate of 257 s^−1^ in the uniform channel (no extensional component), the mean CoA of COL1 fibers was 0.49±0.1 (not aligned) (Figure 3C, S6). Whereas the mean CoA was 0.64±0.11 in segment e of the segmented channel 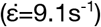 (p<0.05) (**Figure 3C**). These data suggest that the extensional contribution of the flow influenced the alignment in the segmented channel.

### 2.3 Extensional flow promotes fiber alignment over millimeter scales

In the previous experiments, the CoA was measured close to the constriction region **(Figure 3)**. To assess the distance over which the alignment extended into the segment, we imaged COL1 fibers in 3 ROIs spaced 1mm apart within each 5 mm segment **(Figure 4).** We found that the fiber alignment was uniform over each 5 mm segment, with a maximum standard deviation of 0.04 CoA units. The alignment was also consistent through the thickness of the channel **(Movie S1).** These data show that the fiber alignment promoted by the local extensional flow at the constriction extended across the 5mm segment. This rather unexpected finding may be explained by calculating the Weissenberg number 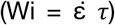, where 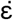 is the extensional strain rate and *τ* is the characteristic molecular relaxation time^52^. Extensional flows are known to stretch and orient molecules^41,53^, and in conditions where W_i_ > 1, molecular orientation effects are favored. In our segmented channel, we found that Wi <1 in the first two constrictions, Wi > 1 in constriction c-d and Wi = 9.1 in constriction d-e (calculations shown in S7). Wi > 1 (segment d) corresponded to the onset of alignment in the segmented channel, suggesting that extensional-flow oriented the self-assembling COL1 fibers in the direction of flow and promoted long-range alignment. Thus, by simply changing the geometry of the microchannel, we could introduce local extensional flow that resulted in uniform fiber alignment across a 5 mm segment, with a CoA that varied as a function of the extensional strain rate at the constriction.

**Figure 4:**
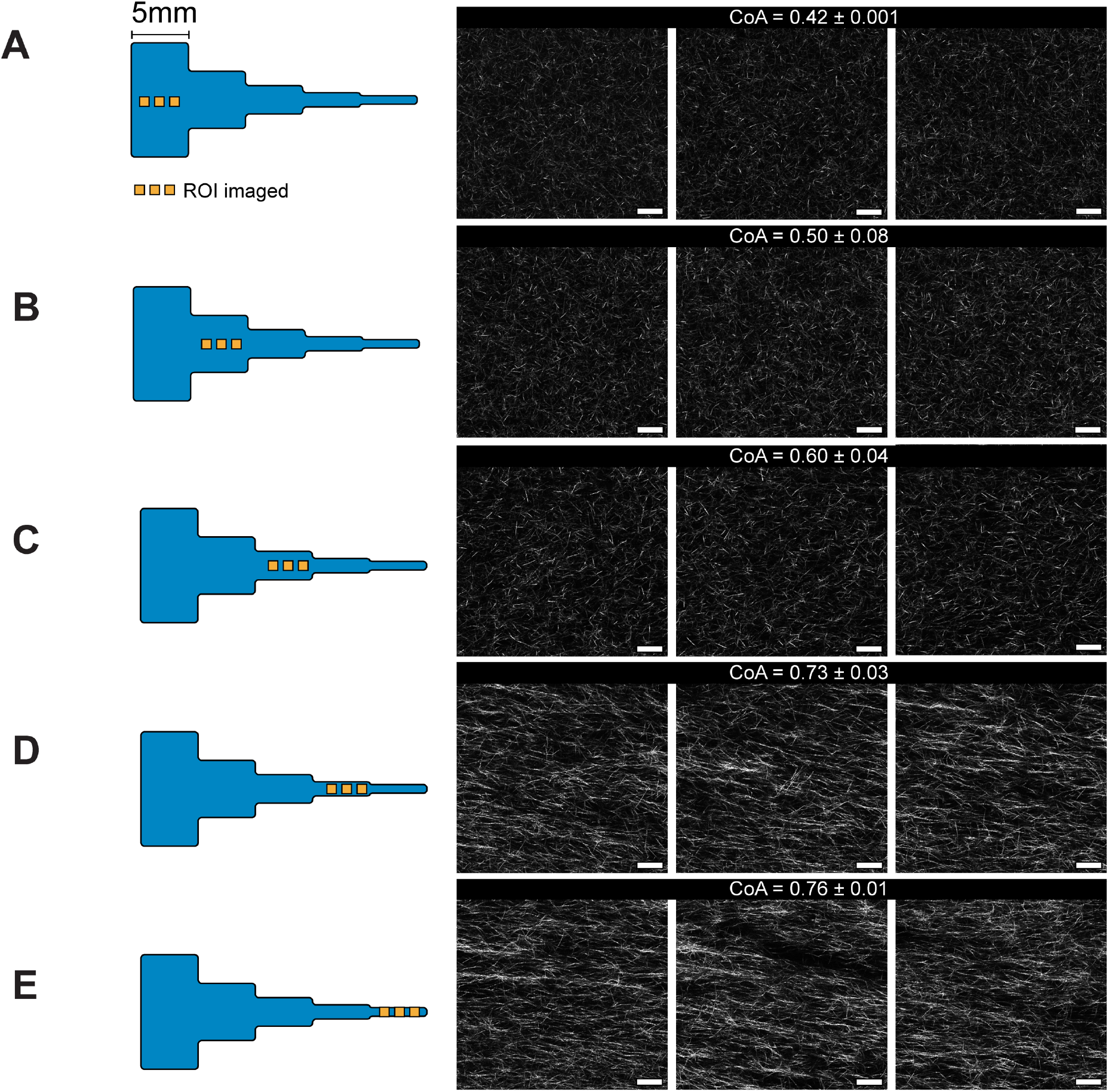
Figure showing CRM images of uniform COL1 fiber alignment across the 5mm segments in the channel. 3 images from each segment **A-E** are shown, spaced 1mm apart. Each image covers an area of 221×221 μm. The mean CoA±SD are shown for each segment, with a maximum deviation of 0.08 units. We can see the onset of fiber alignment in segment **C** that corresponds to W_i_>1 in the constriction b-c. The images are along the flow direction of the neutralized COL1 solution. Scale bar = 25μm.

### 2.4 Cell culture on aligned 3D matrices using functional modules

Having established a relationship between extensional flows and COL1 alignment using a segmented channel, we focused on the practical consideration of how the microengineered 3D COL1 gels could be accessed. Conventional microfluidic systems are permanently sealed, and introducing cells to 3D environments requires that the cells are premixed in the hydrogel precursor solution, or more complex multichannel approaches are needed to deliver media or cells to a hydrogel^31,54,55^. Permanently sealed designs also limit the ability to add new experimental functionalities to an existing device. Here, we take a modular approach that builds on our previous work^56^, where we used magnetic latching to allow our microfluidic device to be easily assembled, disassembled, and combined with specialized modules throughout the experimental workflow.

Figure 5A (1-3) shows the modular platform, consisting of a laser-cut PMMA base with an open central region and holes to accept magnets. The base was mounted onto a glutaraldehyde functionalized glass coverslip. A razor-cut PDMS channel “cut-out” with an outer footprint designed to fit into the open region in the base plate was placed onto the coverslip. A bovine serum albumin (BSA) treated PDMS block containing access ports (referred to hereafter as a “cover”) was placed over the cut-out to form a channel. COL1 solution was injected into the channel and allowed to self-assemble as previously described. After the gel formed, the PDMS cover was peeled away from the cut-out to reveal the aligned COL1 that was confined within the razor-cut features.

**Figure 5:**
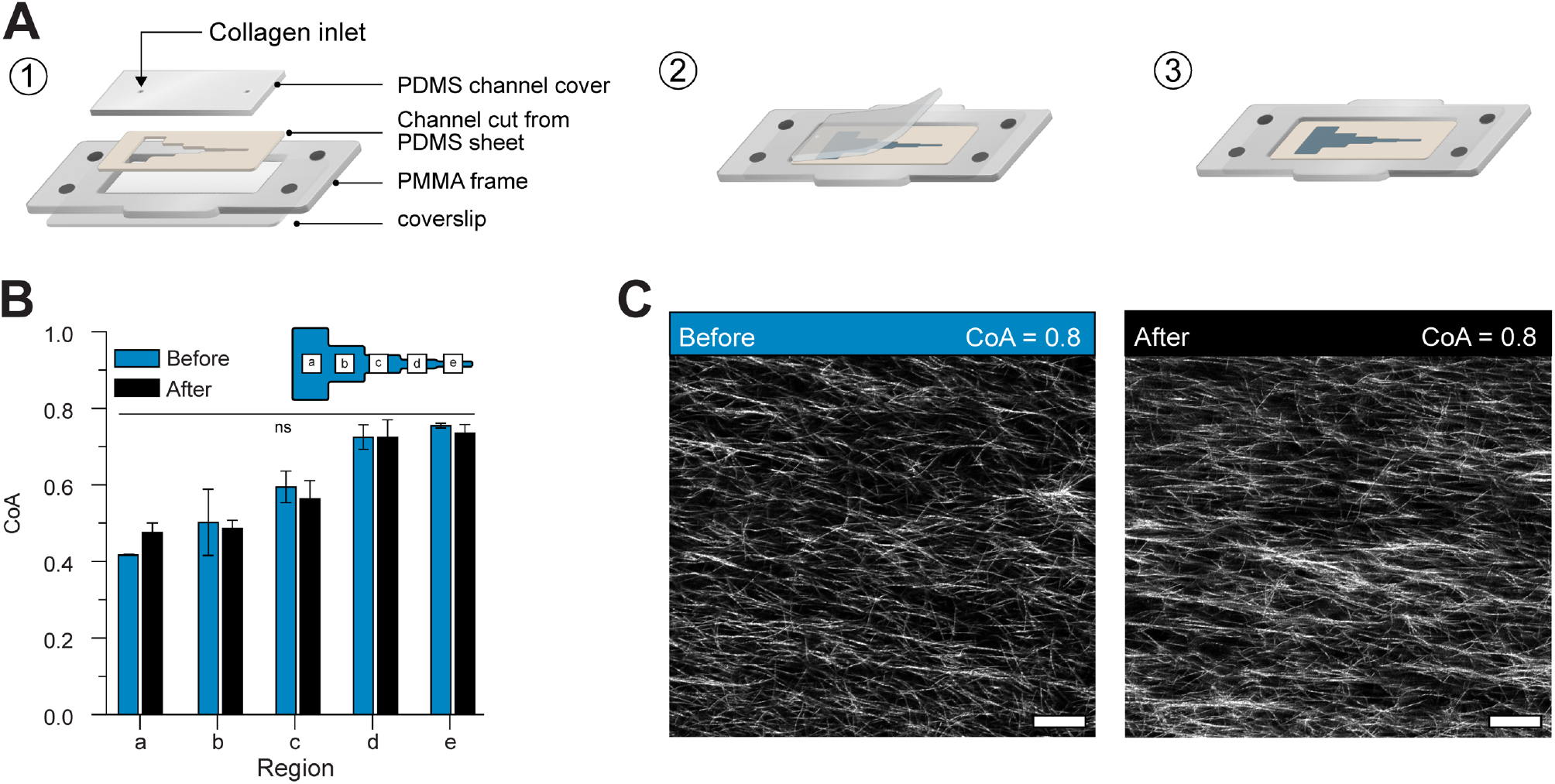
**(A)** Schematic representation of the 2-piece channel fabrication and the modular base. (1) schematic shows the assembly of the channel and the modular base, (2) Schematic illustration of exposing the COL1 matrix in the channel after self-assembly. The cover can easily be removed to reveal the COL1. (3) Direct access to the COL1 matrix after cover peel off. **(B)** Plot shows the COA in each segment of the channel before and after peeling off the cover. No significant difference was observed in the COA. **(C)** Representative images of COL1 fibers in the same region before and after peeling off the cover. Data presented as mean ± SD, n=3, ns=not significant. Scale bar = 25μm

We measured the CoA before and after the process to confirm that the cover removal did not alter the fiber alignment (Figure 5B). Our results showed that the change in CoA was not statistically significant in any of the segments (a-e). Representative images of aligned fibers before and after removing the cover are shown in (Figure 5C). Our cover removal process allowed direct access to the COL1 without the need for auxiliary channels. In addition, the magnetic base allowed the simple “plug and play” addition of specialized functional modules. The magnetic latching process is reversible, quick, and can allow us to add experimental capabilities on-demand without disturbing the underlying COL1 structure. As a demonstration, we first used the modular approach to seed cells and showed differences in endothelial cell alignment on COL1 fibers of high CoA and low CoA.

### 2.5 Positioning endothelial cells using magnetically coupled module

Accurately positioning cells in a large open well is difficult. To position cells on specific segments of the exposed matrix, we first fabricated a module that served as a well to seed cells and held media during the culture process (Figure 6A). The module was constructed by laminating two PMMA pieces with pressure-sensitive adhesive. Integrated magnets allowed the module to be reversibly coupled to the base plate surrounding the two-piece channel (i.e., cutout and cover). After the gel is formed, the cover is removed, and the module fits into the cut-out of the magnetic base and interfaced with the channel and the exposed COL1 matrix, forming a liquid-tight seal against the channel cut-out to hold media or other reagents without leaking. We magnetically coupled the module to the frame and seeded human umbilical vein endothelial cells (HUVECs) in the aligned and unaligned regions of the segmented channel to demonstrate that the cells responded to fiber alignment (Figure 6B). Cells cultured on aligned COL1 (CoA=0.8) displayed an elongated morphology with actin fibers oriented in the direction of COL1 fibers. In contrast, cells interacting with the randomly organized COL1 (CoA=0.3) display random alignment. Using a modular approach, we spatially patterned cell populations and demonstrated cell response to aligned COL1 fibers.

**Figure 6:**
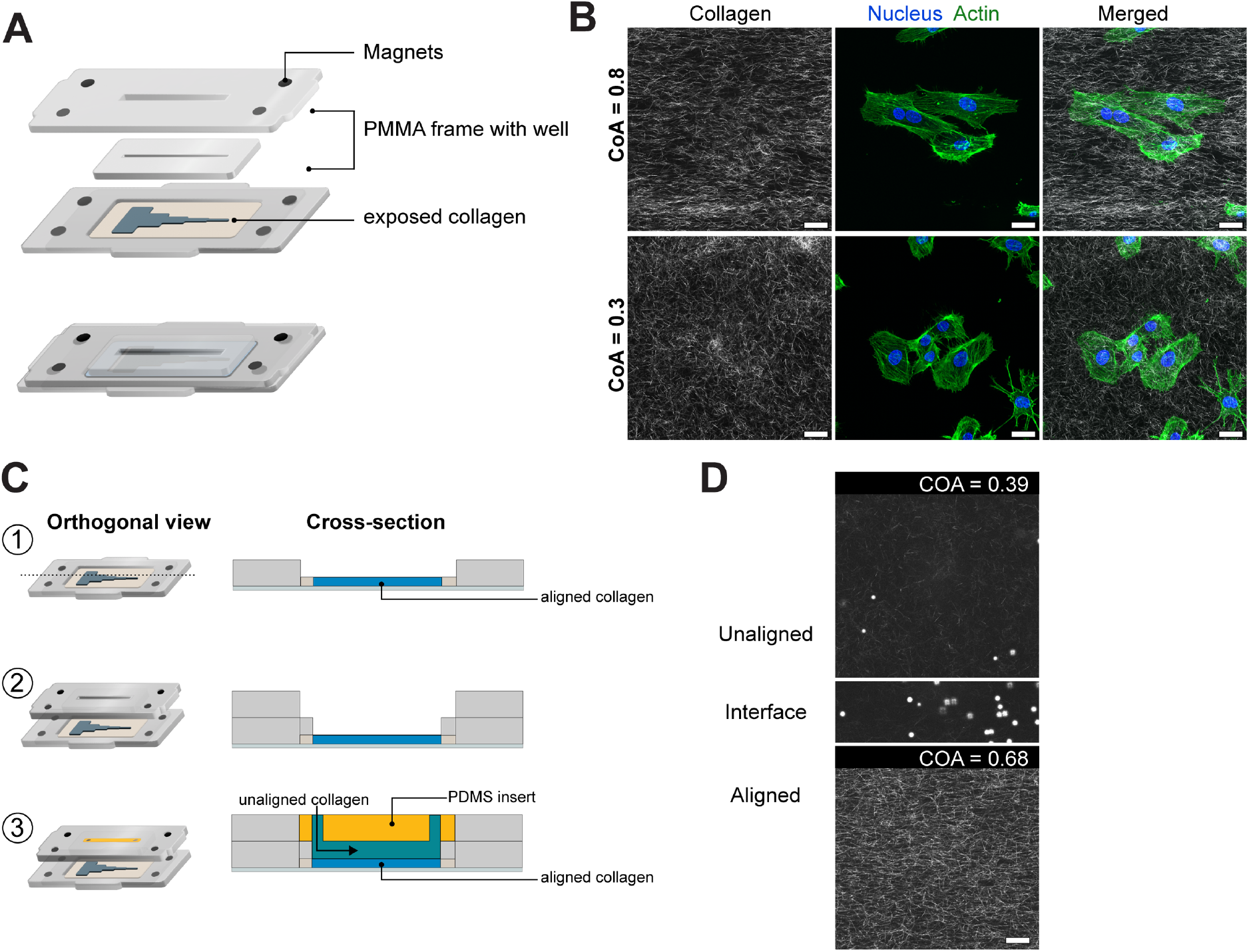
**(A)** Schematic showing the construction and attachment of a cell culture module via magnets on to the aligned COL1 after channel cover removal. The module comprises of a PMMA frame with magnets that enable magnetic coupling to the base. A fully assembled view of the module, after attaching to the base is shown. Laser-cut wells in the module hold cell media. **(B)** HUVECs stained for actin and DAPI after 3 hours of culture on a COL1 matrix in the segmented channel. Top panels show cells in a high CoA (0.8) region, and lower panels show cells in a low CoA (0.3) region of the same hydrogel. Cells are aligned in the aligned region as compared to their morphology in the randomly organized region. **(C)** Module to demonstrate layer-by-layer fabrication of other ECM layers. (1) Collagen exposed after cover removal. (2) Addition of cell culture module. (3) A 2.5mm thick BSA passivated PDMS plug is inserted into the upper half of the 5mm tall well in the fluidic module and Collagen is injected into the 2.5mm chamber below as shown in the cross-section view. **(D)** CRM images of aligned COL1, an interface as marked by 5μm beads, and the top layer of unaligned COL1. Scale bar = 25μm

### 2.6 Layer-by-layer fabrication of COL1 hydrogels

Multi-layered hydrogels can be used to replicate ordered tissue structure^57^, position cell populations relative to other cell types^58,59^, and provide cell-specific niches in a 3D hydrogel. The ability to combine controlled fiber alignment and cell patterning with layer-by-layer biofabrication of 3D hydrogels in a fluidic system can be used to replicate the diverse, ordered microenvironments found in vivo. Here, we used a functional module to fabricate a layer of COL1 with randomly oriented fibers over an existing aligned COL1 substrate (Figure 6C, 1-3). After the self-assembly of the aligned COL1, we removed the channel cover to directly access the microengineered COL1. The cell culture module was magnetically attached, and a BSA passivated PDMS block was inserted into the module, as seen in Figure 6C-3. The PDMS block converted the 5mm deep open well of the cell culture module into a 2.5mm tall channel. Collagen was mixed with polystyrene beads to aid visualization of the layers and injected at a flow rate of 50μL min^−1^ to avoid damaging the existing aligned hydrogel in the segmented channel below. To visualize the different COL1 layers, we imaged the layered construct through its thickness (Figure 6D). The z-stack images revealed a sharp interface between the different COL1 layers, as indicated by the settled polystyrene beads between the aligned and the beads dispersed in the randomly organized COL1 layer. Additionally, the alignment of the pre-existing COL1 in the segmented channel was not affected by the injection of the new layer. We were also able to remove the passivated PDMS cover to expose the randomly oriented COL1 hydrogel layer. This demonstration highlights the ability to fabricate multiple layers of ECM materials within a fluidic device and shows that the addition of new layers did not disturb the microstructure of underlying layers.

## 3. Discussion

### 3.1 The role of extensional flow in COL1 fiber alignment

The use of shear flow is a popular and simple approach to create fiber alignment in thin COL1 gels within a microfluidic channel^26–28^. Most studies that describe shear flows to align COL1 fibers report use acid extracted telocollagen. Telocollagen exhibits faster self-assembly dynamics than atelocollagen which makes it difficult to use in applications where the COL1 solution needs to be manipulated for extended periods of time, such as in microfluidic injections and bioprinting. Recently, the use of bovine-derived atelocollagen has gained popularity in 3D cell culture and as a bioink^45,60–62^. We observed that using shear flows did not induce fibers alignment in thicker hydrogels made from atelo collagen over a wide range of shear rates (50-1000s^−1^). To advance 3D COL1 microengineering capabilities, we tested the hypothesis that the addition of extensional flow promotes alignment in thicker gels. We used a segmented channel with a constant height of 130μm that was suitable for 3D studies^63^ and measured localized extensional flows at each constriction using PIV. We found that the local extensional flow created domains of aligned fibers that were uniformly aligned across each 5 mm segment, and the alignment was a function of the extensional strain rate. This trend was also observed in thicker gels (250 μm), suggesting the possibility of using this process to align thicker hydrogels. The onset of fiber alignment in segment (d) (Figure 3) corresponded to the local extensional flow at the constriction where the Weissenberg number 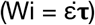 was greater than 1, indicating the importance of the extensional strain rate compared to the relaxation time of the polymer chain (Table S6). To determine whether a single constriction in a channel would align fibers, we flowed COL1 solutions through a channel that contained only one constriction with flow characteristics that corresponded to Wi > 1. The fibers did not align in this single constriction channel configuration (Figure S9), suggesting that repeated exposure to increasing extensional flow is needed to create aligned fibers. This finding is consistent with other reports that observed hysteresis in the conformation of polymer molecules when the molecules were subjected to multiple instances of extensional strain^64^. Thus, our work builds upon recent work that highlights the importance of extensional strain as a driving force in COL1 fiber alignment^12,44^ and shows that different domains of fiber alignment can be microengineered by changing the length of the segments in the channel.

The fiber alignment in our platform can be partially due to the repeated exposure of self-assembling fibrils to extensional strain that may induce fibril stretching and alignment, which further results in the directional growth of the fibril. We hypothesize that CoA values >0.8 can be achieved through tuning COL1 self-assembly rates by using telocollagen instead of atelocollagen, combining COL1 with other biomolecules (e.g., fibronectin, elastin, or hyaluronic acid), increasing the solution pH, or increasing the self-assembly temperature^65–67^. Further optimization of the extensional flows and modifications to the channel geometry, such as the constriction widths, constriction shape, and segment length, may also provide improved alignment. Atelocollagen is less stiff than telocollagen of the same concentration, and the storage modulus of the 2.5 mg mL^−1^ hydrogel was 45Pa (Figure S8). The stiffness of the COL1 matrix can be tuned using similar approaches or by adding exogenous cross-linking agents after the gel cover is removed. We expect that modifying the channel design to generate a desired extensional flow, coupled with control over the COL1 self-assembly conditions, will enable a high degree of control over the fiber alignment in a COL1 matrix that can be rapidly implemented.

### 3.2 Functional Modules and Layer-by-Layer Biofabrication

Since typical microfluidic devices are completely sealed systems, accessing a 3D hydrogel within a microfluidic channel is challenging because loading a hydrogel into the channel effectively blocks fluid delivery ports. Microfluidic devices with auxiliary channels to deliver media and cells into the hydrogel are used to overcome this challenge, but they add complexity to the fabrication process and experimental use. We overcame this challenge by developing a simple two-piece channel that consisted of a channel cut-out and a passivated cover for the channel that was placed on a functionalized glass substrate to enable COL1 adhesion. The cover was used to reversibly seal the channel to inject COL1 into the channel and was removed after COL1 self-assembly to reveal the hydrogel on the glass substrate. The chemical functionalization is standard and ensures that the cover removal did not alter the fiber alignment. We added capabilities for cell patterning and layer-by-layer fabrication by integrating the microfluidic channel into a reversible sealed, easily accessible modular device (SEAM)^56^. The SEAM allowed us to magnetically place functional modules on the exposed COL1 hydrogel for patterning cells and introducing additional ECM layers. The ease of fabricating each component of the device can greatly increase the accessibility of our platform, and additional modules can be designed to advance experimental functionalities.

Typically, the fabrication of microfluidic platforms is carried out using soft-lithography, 3D printing, and in some cases, injection molding. These processes can be time-consuming and require engineering expertise to implement. In our approach, we were able to fabricate both channels and modular platforms using consumer desktop equipment that was well-suited for rapid prototyping. The channel cut-out was fabricated using a desktop razor cutter which reduced the time required for prototyping new channel designs to several minutes, compared to several hours in soft lithography. Similarly, the laser-cut modules were rapidly prototyped without the need for injection molding or milling. Our modular approach offers the potential to customize the experimental platform by attaching additional modules on the microengineered COL1 matrix to pattern cells, perfuse the matrix, and deliver reagents.

Using the modular platform, we have increased the flexibility of adding cell populations at any time in an experiment and patterning multiple cell populations across the segments. The 5 mm long segments with defined alignment are also suitable to investigate the response of cell populations in different microenvironments on the same device. Since the addition of cells is decoupled from collagen preparation, the researcher is now free to manipulate the fibrous microstructure such as pore size, fiber diameter, and stiffness of the COL1 using established methods such as solution preincubation, modifying the pH and ionic strength, and changing the self-assembly temperature ^65–67^. In contrast, premixing cells into a COL1 precursor solution limits the ability to manipulate the self-assembly conditions in COL1 solutions.

The modular platform can also be used to build layered ECM hydrogels with independent control over the composition and mechanical properties of each layer. Layer-by-layer fabrication provides a pathway to generate in vitro models of hierarchal tissues. For example, the cartilage ECM is composed of ordered layers of COL1 rich tissues^68^, and cardiac tissue^69^ features distinct layers of COL1 through the thickness of the tissue with defined degrees of alignment. The layer-by-layer approach can also be used to generate hydrogels with cell-specific layers for 3D coculture in a microfluidic environment. These findings show that extensional flows can be used to microengineer aligned COL1 matrices in a relatively simple manner. Our techniques can also be used to generate alignment in blends of COL1 and other ECM materials such as hyaluronic acid, fibronectin, and fibrin to accurately replicate the composition of the ECM in vitro. We also anticipate that the magnetically coupled modular platform will ease the incorporation of 3D matrices in microfluidic devices while providing new functionalities to fabricate 3D ECM constructs.

## 4. Conclusion

In this work, we found that the addition of local extensional flow using a segmented microfluidic channel design induced COL1 alignment in thick hydrogels. The fiber alignment extended across 5 millimeter segments, and the CoA could be tuned as a function of the applied extensional strain rate within each segment. To access specific aligned domains within each channel segment, we demonstrated a reversibly sealed channel design that allowed for direct access to the self-assembled hydrogel. Using magnetic coupling, we reversibly placed functional modules on top of the aligned COL1 to pattern cells and deposited new ECM layers. We achieved long-range control over the COL1 fiber alignment, cell seeding, and layer-by-layer ECM fabrication using desktop equipment for fabrication and simple chemical functionalization. We anticipate our findings and approach will help customize the microstructure of fibrous ECM hydrogels to establish more complex physiologically-relevant 3D in vitro models.

## 5. Materials and Methods

### 5.1 Soft Lithography

Microfluidic channels were fabricated using standard soft lithography techniques. Channel geometry was designed in Adobe Illustrator (Adobe Inc, CA, USA), and a high resolution, darkfield photomask was printed for lithographically defining a mold (CAD/ART services, OR, USA). A negative photoresist (SU-8 3050, Kayaku Advanced Materials, MA, USA) was spun onto a 4” silicon wafer (UniversityWafer Inc, MA, USA) to a thickness of 130μm following instructions in the manufacturer supplied datasheet. The spin-coated wafer was soft-baked at 90°C for 45 min to evaporate excess solvent and exposed to a 250mJ cm^−2^ dose of 365nm UV to polymerize the resist. The resist was baked at 90°C for 10 min to promote polymerization and developed in SU-8 developer for 15 min, followed by washing in isopropyl alcohol. A laser-cut PMMA frame was placed around the silicon wafer using double-sided pressure-sensitive adhesive (MP468, 3M, MN, USA) to hold PDMS during molding. The mold for the segmented channel was machined from aluminum (RIT Machine Shop) and polished to remove surface aberrations.

To fabricate channels, a PDMS (Sylgard 184, Dow Chemical Company, MI, USA) prepolymer base was mixed with the curing agent at a base to agent ratio of 10:1. The mixture was degassed in a vacuum chamber until no air bubbles were visible in the mixture. The mixed PDMS was poured into the mold and allowed to cure for 10 min at 140°C. After curing, the molded PDMS was gently removed from the mold with tweezers. Inlet and outlet holes were punched using a 1mm biopsy punch (504530, World Precision Instruments, FL, USA). Channels were cleaned by ultrasonication in isopropanol (VWR, PA, USA) for 5 min and rinsing in DI water for 30s, before drying under an air stream. The channels were heated at 100°C for 5 min to ensure they were completely dry and stored in a clean petri dish before use.

### 5.2 Collagen neutralization and injection

Bovine derived atelo-COL1 (Nutragen, Advanced BioMatrix, CA, USA) (416μL) was diluted to a final concentration of 2.5mg/mL^−1^using 10X PBS (100μL) (Thermo Fisher Scientific, MA, USA), ultrapure water (430μL), and 0.1M NaOH (VWR, PA, USA). The volume of NaOH was adjusted to ensure a pH between 8.8-9.0 to increase the rate of fibril formation. pH measurements were carried out on a 25μL aliquot of the prepared COL1 solution, using a pH probe (Orion Star A211, Thermo Fisher Scientific, USA). All reagents were placed on ice and COL1 was injected within 10 min of neutralization.

Collagen injection was carried out in a sterile biosafety cabinet. Collagen solution was loaded into a 1mL syringe (10148-330, VWR, PA, USA) and the syringe was gently tapped to remove air bubbles. The syringe was loaded into a syringe pump (New Era Pump Systems, NY, USA), for controlling the flow rate. A 20-gauge dispensing elbow needle (Grainger Industrial Supply, IL, USA) was attached to the syringe via a luer-lock and used to inject COL1 at specified flow rates into the channels. A minimum volume of 100μL was injected through the channel to minimize effects of the air-liquid interface. Channels were placed in a petri-dish with a damp lint-free wipe to maintain local humidity and prevent evaporation and transferred to a 37°C incubator to induce COL1 self-assembly.

### 5.3 Particle Image Velocimetry

Fluid velocity in the channels was measured using particle image velocimetry carried out using a pulsed laser within a Nikon Eclipse TE2000-S microscope (Nikon Corporation, Japan). 2.5mg/mL Collagen (Nutragen, Advanced Biomatrix, CA, USA) was mixed with 5μm Fluorescent polystyrene latex particles (Magsphere, CA, USA). The COL1 solution was injected at a flow rate of 50μL min^−1^ into the microchannels using a syringe pump (New Era Syringe Pumps, NY, USA). All measurements were conducted in the centerline of the microchannel. For each velocity measurement, 10 pairs of images were captured at a given location and the time interval between images in each pair was set at 300 μs. 3 repeated measures were recorded for each experimental replicate and 3 replicates were measured to calculate mean velocity. All data capturing and analysis were performed in an ROI of 700×700μm using TSI Insight 4G software (TSI, MN, USA). The extensional strain rate was calculated as the change in flow velocity over 700μm.

### 5.4 Rheometry

Rheological properties of the COL1 hydrogels were measured using a Discovery Hybrid Rheometer (HR-2, TA instruments, DE, USA), equipped with a Peltier temperature-controlled stage, 40mm cone geometry with a cone angle of 1°, 40mm diameter and a truncation of 25μm. The geometry was used with a solvent trap to minimize the loss of water content from the COL1 hydrogel during testing. A fresh COL1 hydrogel was prepared at pH 9 as described above, and 330μLwase pipetted on to the Peltier stage at 1°C. The geometry was lowered to a trim gap of 75μm, and any excess COL1 was wiped away before lowering the geometry to a working gap of 25μm. To measure the properties of the neutralized solution as it is kept on ice, the temperature of the stage was maintained at 1°C, a strain of 10% was applied, at 1 frequency of 1Hz for 20 minutes. To measure the viscoelastic properties of the fully polymerized hydrogel, the COL1 was loaded onto the stage at 1°C. The stage was heated to 37°C immediately after starting the experiment and data was recorded for 30 minutes to ensure that the COL1 fully polymerized and the values of storage and loss modulus reached a plateau. To prevent artifacts from COL1 preincubation and potential fibril nucleation, a fresh sample was prepared for each measurement.

### 5.5 PDMS cover passivation

To prevent COL1 from sticking to the PDMS cover, the covers were passivated using BSA. PDMS channel covers were fabricated by casting a 10:1 mix of PDMS (Sylgard 184, Dow Chemical Company, MI, USA) base and curing agent in a PMMA mold. The mold dimensions were 34×14mm. Inlet and outlet ports were cored in the channel using a 1mm biopsy punch (World Precision Instruments, FL, USA). A laser-cut guide was used to help position the inlet and outlet ports. The covers were then cleaned by sonicating in ethanol (VWR, PA, USA) for 5 min, rinsed with DI water for 30sm dried under an air stream, and heated to 100°C for 5 minutes to ensure they were completely dry. The covers were placed in a UV sterilizing chamber for 1 minute to sterilize the surface and immediately transferred to a biosafety cabinet. 350μL of a 40mg/mL^−1^ solution of bovine serum albumin (BSA) (AAJ64100, VWR, PA, USA) in PBS (Thermo Fisher, MA, USA) was pipetted onto each PDMS cover and spread on the surface using a P1000 pipette tip. The covers were kept in a 4°C fridge overnight after which the BSA was aspirated. Thee covers were rinsed thrice with sterile PBS and allowed to dry in the BSC before being placed on the razor-cut PDMS channels.

### 5.6 Coverslip functionalization

24×55mm Glass coverslips (Globe Scientific, NJ, USA) were functionalized with glutaraldehyde (A17876, Alfa Aesar, MA, USA) to promote COL1 attachment to the coverslip via amine-aldehyde-amine coupling. Coverslips were functionalized as described by Syga et al ^70^. Briefly, coverslips were sonicated in 70% ethanol for 5 min followed a rinse in DI water for 1 minute and then dried using pressurized air. The clean coverslips were heated at 100°C for 5 min to ensure that they were completely dry.

To attach amine groups on the surface of the coverslips, the glass surface was first activated by exposing it to a corona discharge wand for 1 min (BD-20AC, Electro-Technic Products, IL, USA). The corona-discharge treated coverslips were immediately dipped in a solution of 2% aminopropyltriethoxysilane (APTES) (80370, Thermo Scientific, MA, USA) in acetone (BDH1101, VWR, PA, USA) for 10 seconds, rinsed with acetone for 30s, and dried under an air stream. A 5% aqueous glutaraldehyde solution was prepared and pipetted over the silanized coverslips. After 30 min, the coverslips were rinsed with DI water for 1 minute, and dried using air. The prepared coverslips were then stored for use up to 3 days in clean petri dishes.

### 5.7 Razor cutting segmented channels

To razor-cut the segmented channels, A 6”x6” PDMS sheet of required thickness (87315K62, McMaster-Carr, IL, USA) was mounted onto an adhesive plastic carrier for easy handling. The carrier sheet was fed into a desktop plotter (Graphtec CE LITE-50, Graphtec, Yokohama, Japan) with an XY plotting resolution of 20μm. The channels were cut at a speed of 1cms^−1^ with the blade depth set to 300μm. After cutting, channels were gently removed from the adhesive carrier sheet using fine-tipped tweezers and sonicated in ethanol (VWR, PA, USA) for 5 min, rinsed in DI water for 30s, and heated at 100°C for 5 min to dry. Channel cut-outs were immediately transferred to a glutaraldehyde treated coverslip after drying.

### 5.8 Modular base fabrication

To implement modular functionality into our platform, we adapted previously described techniques for the fabrication of reversibly sealed, easily accessible modular microfluidic (SEAM) platform^56^. The base was laser-cut using a desktop laser cutter (H-series 20×12, Full Spectrum, CA, USA) from a PMMA sheet (8589K31, McMaster-Carr, IL, USA) of thickness 3/32” (2.38mm), with a 35×15mm well space to accept the PDMS channel cut-out. Rare-earth magnets (4.75mm) (KJ magnetics, PA, USA) press fit into laser cut holes in the base. The base was attached to the glutaraldehyde coverslip using a 50μm thick double-sided pressure-sensitive adhesive (MP467, 3M, MN, USA), and clean channel cut-outs were gently placed on the coverslips. The assembled device was sterilized in a UV sterilizer for 5 min, before being transferred to a sterile biosafety cabinet.

### 5.9 Cell culture module fabrication

The cell culture module is comprised of 2 laser-cut PMMA pieces (8589K31, McMaster-Carr, IL, USA) that are laminated together using pressure sensitive adhesive. The top layer accepts 4 rare-earth magnets (KJ Magnetics, PA, USA) with the opposite polarity facing towards the magnets in the modular base, to enable magnetic latching. The lower layer is attached to the top layer using a double-sided pressure sensitive adhesive (MP468, 3M, MN, USA), with thickness 130μm. and forms a tight seal against the PDMS channel cut-out. The cell culture area is defined by laser cutting a 24 mm x 1.5 mm well in the lower layer and 24 mm x 3 mm in the top layer.

### 5.10 Layer-by-layer fabrication module

The existing cell-culture module could be converted to enable layer-by-layer fabrication. A PDMS plug measuring 24 × 3 × 2.5 mm (l x wd) was fabricated by curing a PDMS solution of 10:1 base to curing agent in a PMMA mold. The plug was inserted into the cell culture module where it fit into the top half of the cell culture well and transformed the 5mm tall well into a 2.5mm thick channel into which a COL1 solution could be injected.

### 5.11 Cell Culture

After self-assembly, the PDMS channel cover was removed to expose the COL1 in the channel. 750μL of endothelial growth media (EGM-2, CC-3162, Lonza, Basel, Switzerland) was added to the well and COL1 was incubated in media for 24 hours before seeding cells. Human umbilical vein endothelial cells (HUVECs) were obtained from Thermo Fisher Scientific. Cells were cultured in endothelial cell growth medium (EGM-2 CC-3162, Lonza, Basel, Switzerland), supplemented with BulletKit (CC-4176, Lonza, Basel, Switzerland), at 37°C, 95% humidity. Cells were passaged at 70% confluence and used between passages 3-6. For seeding in modular device, cells were enzymatically dissociated from the culture dish using Trypsin (Corning, NY, USA) for 3-5 min and centrifuged at 150G for 5 min, resuspended, and counted using an automated cell counter (Countess II, ThermoFisher, USA). Collagen gels were gently washed twice with media and the cell culture module was magnetically attached to the modular base. Cells were seeded at a density of 10,000 cm^2^ in the cell culture module.

### 5.12 Immunostaining and Imaging

Cells were fixed in 4% paraformaldehyde (Electron Microscopy Sciences, PA, USA) after 3 hours of seeding. Cells were permeabilized using Triton X-100 (Alfa Aesar, MA, USA) for 15 min, and blocked with 4% BSA (AAJ64100, VWR, PA, USA) in PBS. Cells were washed with PBS Tween-20 (PBST) (Thermo Fisher Scientific, MA, USA) after each step. Cells were labelled with phalloidin conjugated to AlexaFluor-488 (A12379, Thermo Fisher Scientific, MA, USA) for 30 min (1:400 dilution), followed by Hoechst 33342 (1:500 dilution) (H3570, Thermo Fisher Scientific, MA, USA) for 10 min to stain actin fibers and cell nuclei respectively. Lastly, cells were washed with PBST for 15 min, followed by a washing with 1X PBS for 5 min before imaging. Imaging was carried out using a Leica SP5 confocal microscope (Leica Microsystems, Germany) using a 40X water objective (HCX PL APO 40x/1.1 W), and optical zoom of 1.75x. Laser line of 488nm was used to visualize actin stained with AlexaFluor-488, and reflectance imaging of COL1. 405nm diode laser was used to visualize Hoechst 33342 labelled nuclei. 15μm thick z-stacks were captured for Hoechst and Phalloidin, with 115 slices per stack. Collagen was imaged 10μm below the cell surface, to visualize the underlying fiber alignment without background from the cells. Stacks captured were of 2μm thickness, with 15 slices in each stack. Image channels were merged and projected using a maximum projection, and a scale bar added using FIJI (NIH, USA). Collagen channels were projected using average projection to reduce noise.

### 5.13 Quantification of fiber alignment

Collagen fiber images z-stacks were flattened using average intensity projection using FIJI. LOCI CT-FIRE, a curvelet transform based package was used to identify individual fibers in the confocal reflectance microscopy (CRM) image^71,72^. A histogram of fiber orientations was made, with intervals of 15°. The number of fibers that were oriented 15° above or below the mode of the orientation histogram was recorded and divided by the total number of detected fibers to get the coefficient of anisotropy (CoA). The CoA ranged from 1 (perfectly aligned) to 0 (complete randomness). CoA > 0.5 was aligned.

### 5.14 Data Analysis

To determine the alignment of COL1 fibers in segmented channels, 3 images were captured in each segment, and 4 independent samples were investigated. In the uniform channels, 3 images were captured in each channel, across 3 independent samples for each shear rate. Data were analysed using GraphPad Prism 9.0 (GraphPad Software, CA, USA). One-way ANOVA was used to determine statistical significance. All data is presented as mean ± SD, N=3 except where mentioned. p<0.05 was considered to be statistically significant.

## Acknowledgements

This work was supported in part by the National Institute of Health (NIH) under award numbers R35GM119623 and R61HL15434. The authors acknowledge Xian Bowles for illustration support and members of the Biological Microsystems Laboratory at RIT for insightful scientific discussions. The content of this publication is solely the responsibility of the authors and does not necessarily represent the official views of the NIH.

---

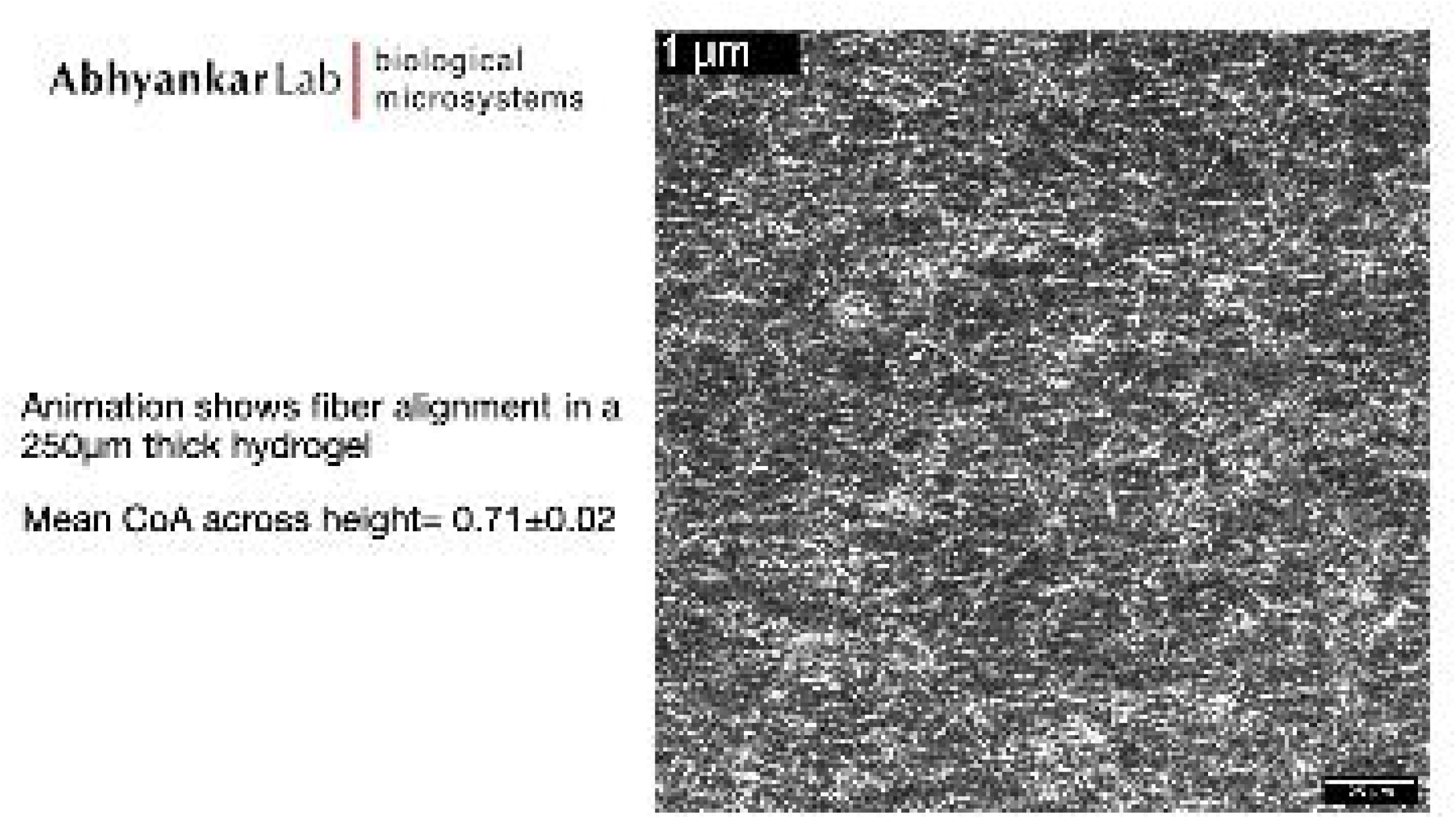

