## supplemental data for "Local Extensional Flows Induce Long-Range Fiber Alignment in 3D Collagen Hydrogels"

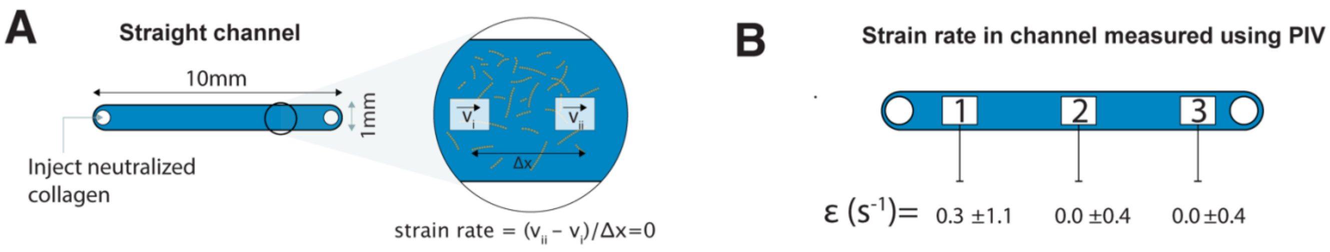


Figure S1: Schematic showing the dimensions of the straight channel with zero-strain condition. (B) Strain rate measured using PIV shows that the extensional strain in the straight channel was ~0 $s^{-1}$

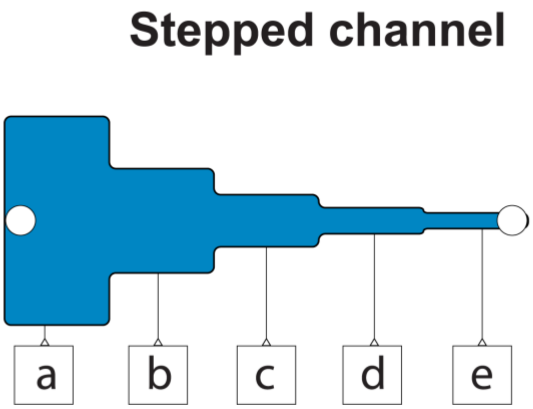


| ROI | strain rate |
|---|---|
| a | 0.16±0.1 |
| b | 0.39±0.6 |
| c | 0.24±0.3 |
| d | 0.26±0.3 |
| e | 0.21±0.5 |

Figure S2: extensional strain rates measured in the center of each segment using PIV show that $\dot{\varepsilon}<0.5$ in the center of all segments. The extensional strain was confirmed to as being localized to the constrictions.

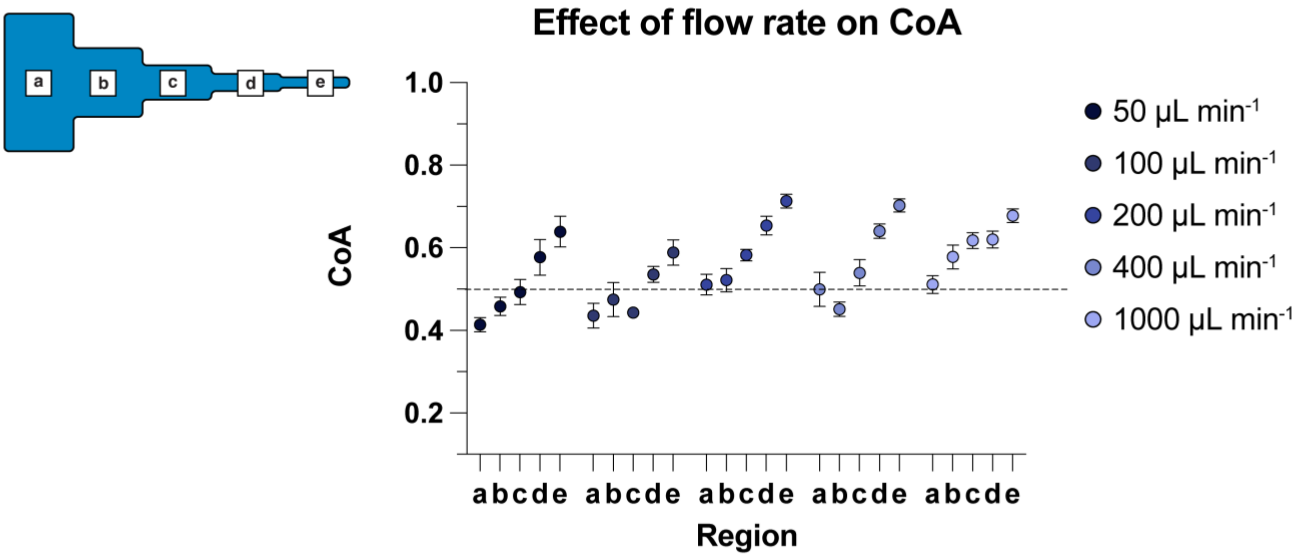


Figure S3: Plot shows the COA of collagen fibers injected at flow rates from 50-1000μL min$^{-1}$ in 130μm thick segmented channels. The fiber alignment increased from segment (a) to segment (e) in all conditions. With increasing flow rate, the maximum collagen alignment increased to a peak alignment of ~0.8.

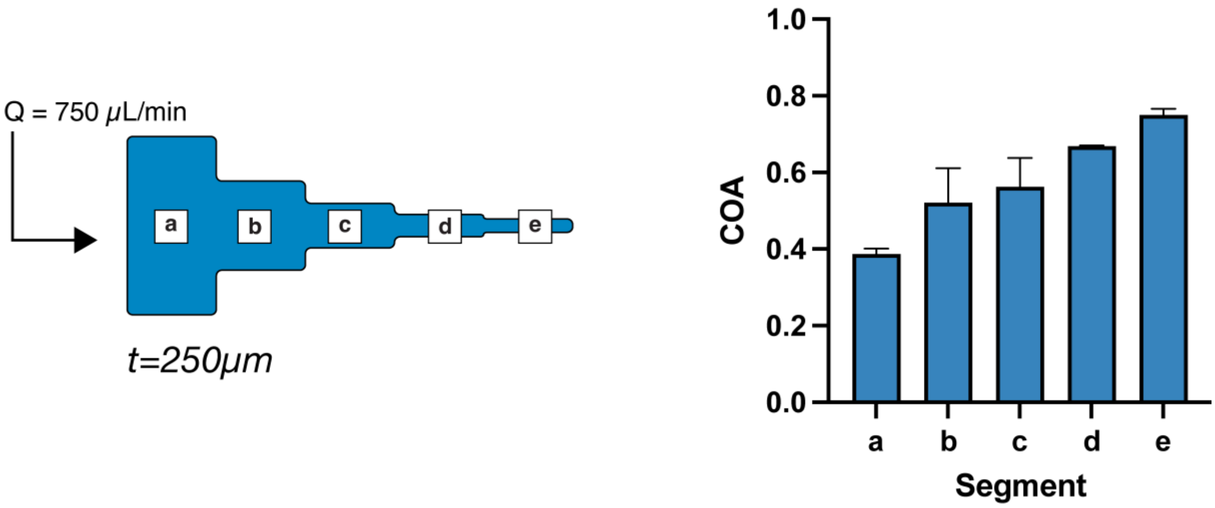


Figure S4: COA measurements of collagen injected at Q=750μL min$^{-1}$ into a 250μm thick segmented channel. Mean COA increased going from segment (a) to (e) with a peak average of 0.75.

S5: Calculated average shear rate in each segment:

| Maximum centerline velocity (PIV) (mm/s) | Shear rate ($s^{-1}$) |
|---|---|
| 1.531 | 23.55 |
| 2.55 | 39.28 |
| 3.98 | 61.23 |
| 8.32 | 128 |
| 16.7 | 257 |

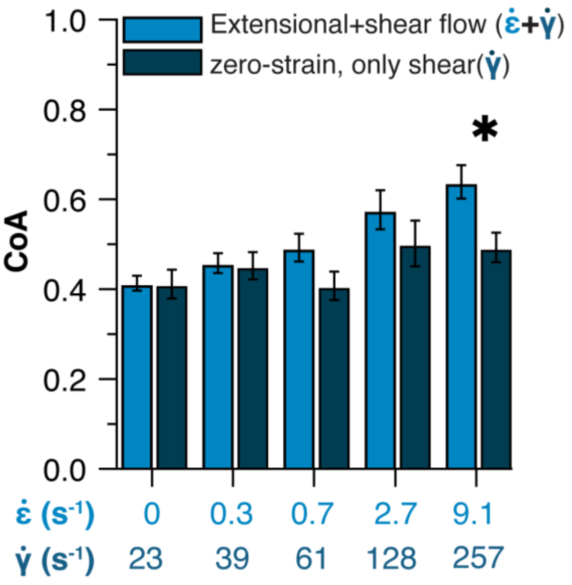


Figure S6: Data showing collagen fiber alignment under extensional and shear flows. The light blue bars show the mean CoA of fibers that were exposed to extensional flow (segmented channel), and the dark blue bars represent the mean CoA of collagen fibers exposed to only shear (straight channels). N=3, Mean±SD, *P<0.05

S7:

To determine the relaxation time, we used the Rouse model, as used in other studies earlier[1]. Briefly, the relaxation time is calculated as:

$$\tau_r = \frac{C_d R^2}{6\pi^2 K_B T}$$

Where, $C_d$ is the drag coefficient of the subunit, calculated as

$$C \quad d = \mu \frac{2\pi L}{ln\frac{L}{d}}$$

and $R^2 = 2 l_p L$, where $\mu$ is the viscosity of the solution, $L$ is the length of the subunit, $l_p$ is the persistence length, and $d$ is the subunit diameter.

In a fibril forming environment, the average collagen fibril length has been reported as $L$= 255.6 $nm$, with a persistence length $lp$=18038±1870 $nm$ [2] .

Using the equations above, the Weissenberg number was calculated to range from 0.30 in the first constriction, to a high of 4.16 in the smallest constriction. we notice that the onset of fiber alignment correlates to $W_i$=1.28

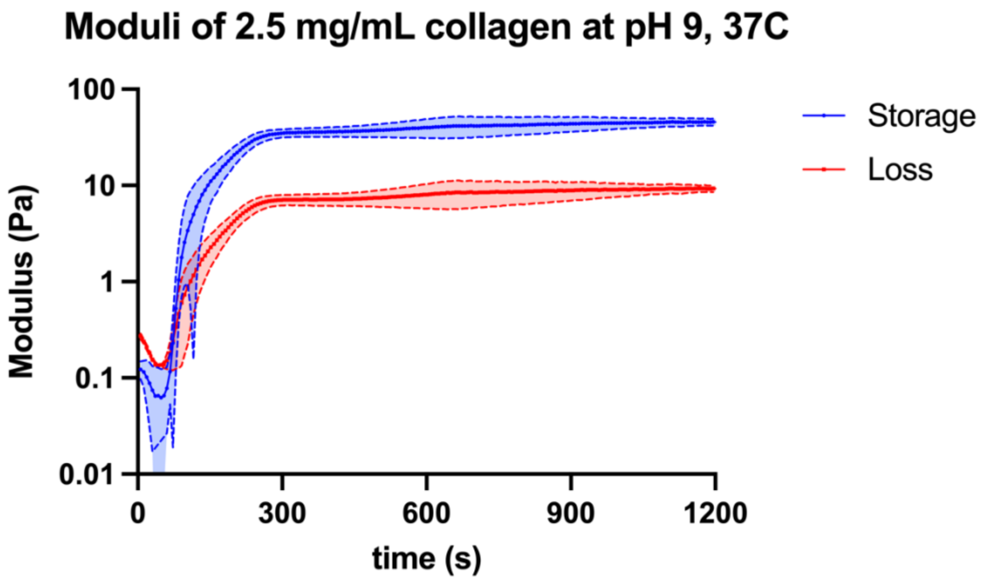


**Figure S8:** Rheometry data showing the increase in storage and loss modulus of the collagen during self-assembly at 37C. Mean storage modulus at the plateau was measured to be 45 Pa. N=3

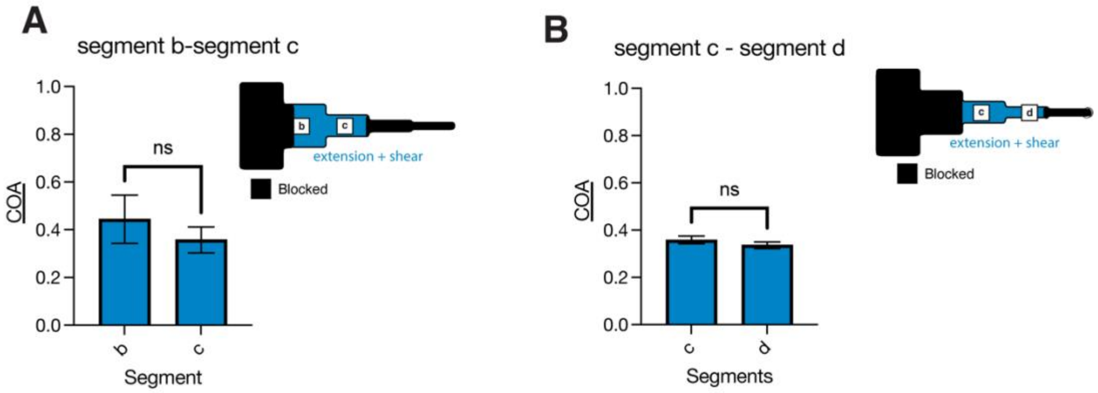


**Figure S9**: Plot shows the COA of collagen fibers that were injected in channels with a single constriction. Constrictions before and after the segments of interest were cut away and (A) the channel was designed to only present constriction between segment b-segment c, corresponding to a strain rate of 0.66 $s^{-1}$. (B) Channel with constriction between segments (c)- segment (d), strain rate = 2.73 $s^{-1}$. ns = not significant
